# Genetic characterization of African swine fever virus in Romania during 2018-2019 outbreak

**DOI:** 10.1101/2019.12.15.876938

**Authors:** Vlad Petrovan, Mihai Turcitu, Lilia Matei, Vlad Constantinescu, Mihaela Zaulet

## Abstract

African swine fever (ASF) is a highly contagious and lethal viral disease of swine with significant socio-economic impact in the developed and developing world. Since its reintroduction in 2007 in the Republic of Georgia, the disease has spread dramatically thorough Europe and Asia. Among the most affected countries in Europe is Romania, which initially reported the disease in 2017 and in 2018-2019 lost about 1 million pigs. There is no molecular characterization of the virus circulating in Romania during that reported period; therefore, the purpose of this study was to provide an initial molecular characterization using samples collected from two farms affected by ASFV. The causative strain belongs to genotype II, and its closest relatives are the strains circulating in Belgium, Russia, and China.

## Introduction

African swine fever (ASF) is a lethal hemorrhagic fever of swine and is the most important foreign animal disease threatening the worldwide agriculture, currently spreading in Europe and Asia. Since there is no vaccine or treatment available, strict biosecurity measures and trade restrictions are implemented during an initial outbreak (Sánchez-Vizcaíno, 2015). ASF is caused by a macrophage-tropic, double stranded (ds) DNA virus with a 170–190 kb genome, currently the only member of the *Asfarviridae* family (Dixon, 2011). The principal routes of disease transmission are direct contact between infected pigs and indirect contact through contaminated feed and food products (EFSA, 2014). Moreover, the virus is maintained in the sylvatic cycle through the soft ticks of genus *Ornithodoros* (Diaz, 2012).

The disease was initially described in Kenya in the early 1900s, causing high morbidity and mortality among domestic pigs (Montgomery, 1921). The first transcontinental spread of African swine fever virus (ASFV) occurred in 1957, in Portugal and Spain (Manso Ribeiro, 1958). As of today, ASF still remains endemic in Africa and on the island of Sardinia, in Italy. Since then, ASFV spread again out of Africa to the Caucasus and subsequently to Eastern Europe, resulting in outbreaks in the Russian Federation and in neighboring countries, including Belarus, Ukraine, Lithuania, Estonia, Poland, Latvia, Czech Republic, Moldova, Romania and Hungary. Recently, ASFV outbreaks have occurred in major swine-producing countries in China, Mongolia, and Vietnam. ASFV has a wide range of genetic variation (24 different genotypes), as shown by the sequencing of the C-terminal end of the major capsid protein p72 and full sequencing of p54. Variation between closely related genotypes is shown by sequencing the central variable region (CVR) of B602L (Bastos, 2004; Achenbach, 2017; Quembo, 2018).

Since the first case in 2017, Romania’s National Sanitary Veterinary and Food Safety Authority (NSVFSA) confirmed more than 1000 ASF cases, including large scale biosecurity facilities and backyard farms (OIE, 2019). Therefore, Romania’s economy had suffered major economic losses among European Union (EU) members due to massive depopulation strategies. However, there is no molecular characterization available to monitor ASFV evolution in Romania.

The purpose of the present study was to perform initial genetic characterization using classical approach of genotyping of p72, p54 and CVR of B602L (Bastos, 2004; Achenbach, 2017; Quembo, 2018).

## Materials and methods

Tissue samples from four infected pigs were collected from two backyard pig farms during 2018-2019 outbreak (between September 2018-February 2019). Samples were collected from one farm in N-W Romania, Bihor county (September 2018) and one farm in S-E Romania from Braila county (February 2019), were diagnosis of ASF was confirmed by the NSVFSA local laboratories. DNA was extracted from tissue homogenates (spleen, tonsil, and lymph nodes) of each animal. Next, we performed PCR analysis using specific primers used for genotyping (Gallardo, 2009). The DNA sequencing reactions were performed as described previously (Zaulet, 2012). Briefly, the reaction products were purified using Wizard® PCR Preps DNA Purification System (Promega, Madison, WI, USA), and the concentration and purity of the products were evaluated by spectrophotometry (Eppendorf BioPhotometer, Hamburg, Germany). The sequencing was performed on a 3130 Genetic Analyzer and the obtained sequences were truncated manually and received GenBank accession numbers (Figure 1). Sequence alignments and phylogenetic analysis were performed using CLC Workbench software v 7.6.3 using a set of reference sequences corresponding to all 24 genotypes of ASFV.

**Figure 1.**
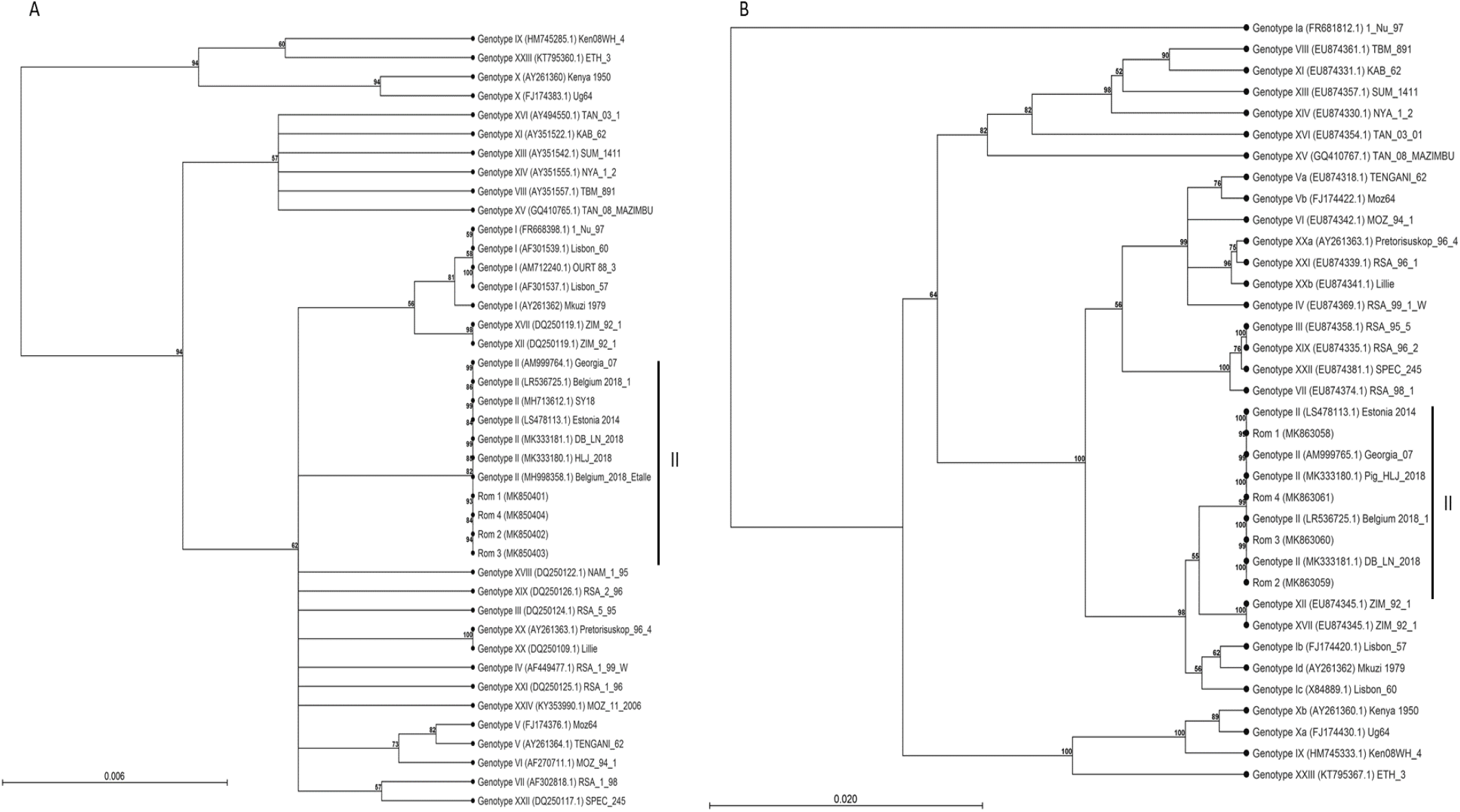
Minimum evolution (ME) phylogenetic trees for major capsid protein gene (p72; panel A) and envelope protein (p54; Panel B) of ASFV isolated during 2018-2019 outbreaks in Romania. Corresponding genotypes are labeled I-XXIV. The strain name and GenBank accession number are indicated. Vertical black lines indicate the genotype from Romanian sequences generated during this study. Scale bar indicates nucleotide substitutions per site. The percentage of replicate trees >50% in which the associated taxa clustered together by bootstrap analysis (1,000 replicates) is shown adjacent to the nodes.

## Results and discussion

Two data sets were generated for the phylogenetic analysis 1) p72 analysis using 41 sequences corresponding to all currently available ASFV genotypes and 2) p54 analysis using 36 sequences corresponding to all the sub-genotypes available for ASFV (excluding genotypes XXII and XXIV, which there were no sequences available for p54). Unweighted pair-group arithmetic average (UPGMA), neighbor joining (NJ), and minimum evolution (ME) phylogenetic tress were constructed using Kimura 2-parameter substitution model, as determined by a model selection analysis used by CLC Workbench v. 7.6.3 (Figure 1, panel A for p72 and panel B for p54). Bootstrap analysis was performed 1,000 times to assess the degree of statistical support for the resulting p72 and p54 trees.

Phylogenetic analysis revealed that the strain currently circulating in Romania belongs to genotype II, and is identical with the ones described in Georgia, Russia, China and Belgium for both p72 and p54. We obtained similar results by sequencing CVR within B602L revealing 100% identity with the isolates currently circulating in Ukraine and Russia (data not shown) (Gallardo, 2014). The results obtained confirm that evolution of ASFV in the Romanian pig farms follows one evolutionary direction. It is important to note that the samples were collected from two different regions during the outbreak season of 2018-2019; therefore, our analysis revealed that the virus did not acquire any additional mutations in the three genes used for genotyping.

However, a further genome-wide genotyping focusing on variable intergenic markers will consolidate our findings and bring more information regarding ASFV evolution in Romania and in Eastern Europe.

